# DRAM1 requires PI(3,5)P_2_ generation by PIKfyve to deliver vesicles and their cargo to endolysosomes

**DOI:** 10.1101/2020.12.15.422832

**Authors:** Michiel van der Vaart, Adrianna Banducci-Karp, Gabriel Forn-Cuní, Philip M. M. Witt, Joost J. Willemse, Salomé Muñoz Sánchez, Rohola Hosseini, Annemarie H. Meijer

**Affiliations:** Animal Sciences, Institute of Biology Leiden, Leiden University, Leiden, 2333CC, The Netherlands

## Abstract

Endolysosomal vesicle trafficking and autophagy are crucial degradative pathways in maintenance of cellular homeostasis. The transmembrane protein DRAM1 is a potential therapeutic target that primarily localises to endolysosomal vesicles and promotes autophagy and vesicle fusion with lysosomes. However, the molecular mechanisms underlying DRAM1-mediated vesicle fusion events remain unclear. Using high-resolution confocal microscopy in the zebrafish model, we show that mCherry-Dram1 labelled vesicles interact and fuse with early endosomes marked by PI(3)P. Following these fusion events, early endosomes mature into late endosomes in a process dependent on the conversion of PI(3)P into PI(3,5)P_2_ by the lipid kinase PIKfyve. Chemical inhibition of PIKfyve reduces the targeting of Dram1 to acidic endolysosomal vesicles, arresting Dram1 in multivesicular bodies, early endosomes, or non-acidified vesicles halted in their fusion with early endosomes. In conclusion, Dram1-mediated vesicle fusion requires the formation of PI(3,5)P_2_ to deliver vesicles and their cargo to the degradative environment of the lysosome.

## Introduction

Endocytic processes are specialised in the uptake of substances from the microenvironment of the cell. Although most of the endocytic cargo is used for cellular sustenance or recycled back to the plasma membrane, a proportion of endocytosed material (e.g. pathogens and remnants of dead cells) is routed towards acidic and hydrolytic lysosomes for its degradation (Huotari and Helenius, 2011). While the endolysosomal system is responsible for degradation of unwanted extracellular material, autophagy performs a similar housekeeping function for the removal of intracellular material. During autophagy, cytoplasmic content is captured in double membraned vesicles and delivered to the endolysosomal system for degradation and recycling (Glick et al., 2010). In this way, autophagy replenishes nutrient levels in times of cellular starvation, and clears the cytoplasm of unwanted elements like protein aggregates, malfunctioning organelles, and intracellular pathogens (Saha et al., 2018). Routing endosomal and autophagosomal content to the degradative environment of the lysosome requires multiple vesicle fusion and maturation steps. Disruptions in these processes can result in serious pathologies, including neurodegenerative diseases, lysosomal storage disorders, infection, and cancer (Cossart and Helenius, 2014; Deretic et al., 2013; Malik et al., 2019; Sun, 2018; Tzeng and Wang, 2016).

DNA damage regulated autophagy modulator 1 (DRAM1) regulates autophagy and endolysosomal fusion events. DRAM1 was first identified as a cellular stress-induced regulator of autophagy and cell death downstream of tumour suppressor protein p53 (Crighton et al., 2006). DRAM1 primarily localises to lysosomes but can also be detected on other organelles of the vesicular trafficking system, including endosomes, autophagosomes, autolysosomes, the Golgi apparatus, and the endoplasmic reticulum (Crighton et al., 2006; Mah et al., 2012). Furthermore, DRAM1 was found to regulate fusion between autophagosomes and lysosomes, a process called autophagic flux (Zhang et al., 2013). When host cells detect pathogenic mycobacteria that cause tuberculosis, *DRAM1* expression is activated downstream of the immunity regulating transcription factor NFκB (van der Vaart et al., 2014). Knockdown or knockout of *dram1* increased susceptibility to mycobacterial infection in zebrafish, identifying Dram1 as a host resistance factor (van der Vaart et al., 2014; Zhang et al., 2020). In support of this finding, overexpression of *dram1* was protective against mycobacterial infection by enhancing autophagic defences against intracellular bacteria and stimulating vesicle fusion events with lysosomes (van der Vaart et al., 2014). Although the effects of DRAM1 activation on autophagy and endolysosomal fusion events are described for several situations, its underlying molecular function in these processes remains unknown.

Endocytic cargo is sorted in early endosomes marked by the GTPase Rab5 (Zerial and McBride, 2001). Early endosomes containing cargo destined for degradation gradually replace Rab5 on their membrane for Rab7, while lowering their luminal pH from values above pH 6 to pH 6.0-4.9 to become late endosomes (Maxfield and Yamashiro, 1987; Zerial and McBride, 2001). During this phase of the maturation process, the outer endosomal membrane starts budding inwards to form intraluminal vesicles (Raiborg et al., 2002; Sachse et al., 2002). The resulting multivesicular bodies are a type of late endosome that also receive cargo destined for degradation by fusing with autophagosomes (Fader and Colombo, 2009). Late endosomes then undergo transient ‘kiss-and-run’ interactions with lysosomes, before eventually undergoing full fusion with lysosomes to reach the endpoint of this degradative pathway at a luminal pH of around 4.5 (Maxfield and Yamashiro, 1987). This process of endolysosomal maturation is extensively reviewed by Huotari & Helenius (Huotari and Helenius, 2011).

The identity of endolysosomal vesicles is in part determined by the presence of phosphoinositide (PI) lipids in their membrane that serve as docking sites for effector proteins (Hammond et al., 2012; Heo et al., 2006; Strahl and Thorner, 2007). Pls can be phosphorylated and dephosphorylated on the hydroxyl groups at the three, four, and five positions of their inositol rings by a range of kinases and phosphatases, generating a total of 7 different Pls in animals (Banerjee and Kane, 2020). Typically, early endosomes are defined by the presence of PI(3)P in their membrane, which is converted into PI(3,5)P_2_ by the lipid kinase PIKfyve (Fab1p in yeast) during maturation into late endosomes (Wallroth and Haucke, 2018). Deletion or inhibition of PIKfyve/Fab1p results in accumulation of enlarged early/late hybrid endosomes that contain few intraluminal vesicles (Cai et al., 2013; Futter et al., 2001; Ikonomov et al., 2003; Jefferies et al., 2008; Odorizzi et al., 1998).

To gain a better understanding of the mechanisms behind DRAM1-mediated vesicle fusion events, we used the zebrafish *in vivo* model to study the potential link between DRAM1 and PI lipids involved in endolysosomal fusion and maturation. By generating a transgenic line that ubiquitously expresses fluorescently-tagged Dram1, we were able to confirm that Dram1 primarily localises to acidic vesicles. High-resolution confocal time-lapse imaging revealed that fluorescently-tagged Dram1 labels dynamic vesicles that display either a globular or tubular morphology. These Dram1-positive vesicles interact and fuse with early endosomes containing PI(3)P in their membranes. Early endosomes that have fused with Dram1-positive vesicles mature into late endosomes as they gradually reduce the presence of PI(3)P lipids in their membranes. Inhibition of PIKfyve, which prevents the formation of PI(3,5)P_2_, reduced the targeting of Dram1 to acidic endolysosomal vesicles (late endosomes and lysosomes). Instead, fluorescently-tagged Dram1 accumulated in multivesicular bodies, early endosomes, and non-acidified vesicles halted in their fusion with early endosomes. Based on these findings, we conclude that Dram1-mediated vesicle fusion is dependent on the formation of PI(3,5)P_2_ by PIKfyve to deliver vesicles and their cargo to the degradative environment of the lysosome.

## Results

### mCherry-Dram1 labels vesicles that interact and fuse with early endosomes

The molecular function of DRAM1 in vesicle trafficking remains largely unknown. To understand the breadth of it’s possible functions, we used the Eukaryotic Linear Motif (ELM)(Kumar et al., 2020) resource to predict functional sites in the human DRAM1 protein (Figure 1A). This analysis confirmed the previously reported presence of 6 transmembrane domains (Crighton et al., 2006), suggesting that DRAM1 is embedded in cellular membranes with parts of the protein exposed to opposite sides of this membrane. Amongst the predicted protein domains, we identified two domains that support a function for DRAM1 in vesicle trafficking. Eps15 homology (EH) domains are generally present in proteins that regulate endocytosis or vesicle trafficking processes (Naslavsky and Caplan, 2005). The autophagy-related protein Atg8 and its homologs LC3 and GABARAP are markers of autophagosomes (Glick et al., 2010). The presence of Atg8 interacting domains therefore suggests that DRAM1 can interact with the autophagy-machinery.

**Figure 1:**
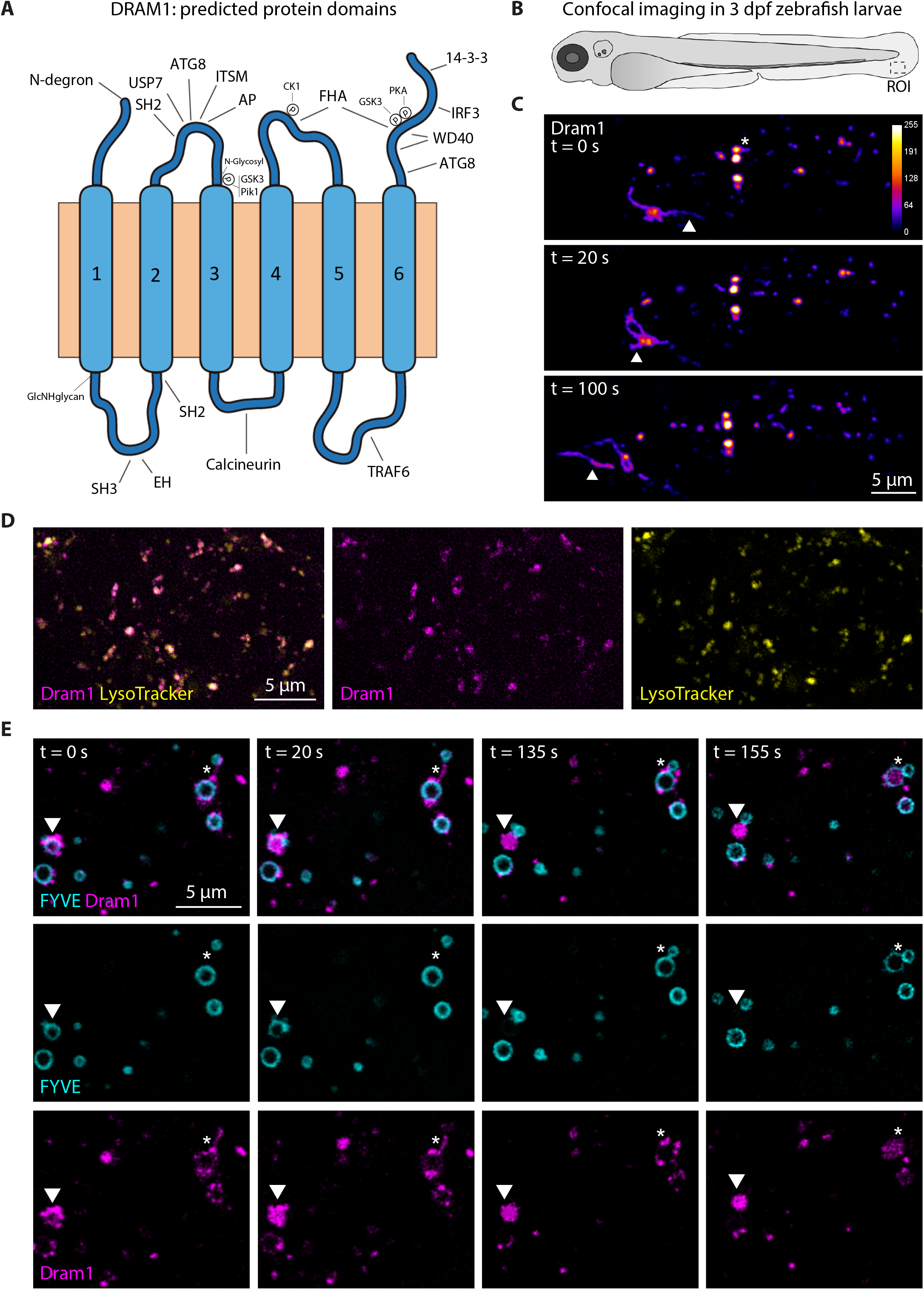
Transmembrane protein Dram1 mainly localises to acidic vesicles and interacts with early endosomes. (A) Schematic representation of protein domains predicted by the Eukaryotic Linear Motif (ELM) resource in the human DRAM1 protein (UniProtKB: Q8N682). (B) Schematic representation of the region of interest (ROI) used for confocal imaging of basal cell layer epithelial cells in the tailfin of 3 days post fertilisation (dpf) zebrafish larvae. (C) Representative stills from time-lapse confocal imaging of mCherry-Dram1. A globular mCherry-Dram1 labelled vesicle displaying a tubular extension is indicated by an asterix (*). A motile tubular mCherry-Dram1 labelled vesicle is indicated by arrowheads (Δ). The intensity calibration bar for the Lookup table (LUT) is displayed in the top panel, ranging from 0 to 255. (D) Representative maximum intensity Z-projection of basal cell layer epithelial cells expressing mCherry-Dram1 and stained with LysoTracker Deep Red. Panels (from left to right) display the merged image, mCherry-Dram1 in magenta, and LysoTracker Deep red in yellow. (E) Representative stills from time-lapse confocal imaging of mCherry-Dram1 and GFP-2xFYVE in basal cell layer epithelial cells. The top panels display the merged image for each time point with mCherry-Dram1 in magenta and GFP-2xFYVE in cyan, while the panels below show GFP-2xFYVE and mCherry-Dram1 seperately. A tether formed by mCherry-Dram1 between two GFP-2xFYVE labelled vesicles is indicated by an asterisk (*). The arrowheads (Δ) indicate a GFP-2xFYVE labelled vesicle that fuses with mCherry-Dram1 (t=0s and t=20s) and subsequently loses the GFP-2xFYVE labelling of its membrane (t=135s and t=155s). Scale bars: 5 μm.

We aimed to study the dynamic localisation of DRAM1 during endolysosomal maturation processes. For this purpose, we used a previously described mCherry-Dram1 construct under control of the ubiquitous beta actin promoter to generate a transgenic zebrafish line fluorescently reporting the subcellular localisation of Dram1 (van der Vaart et al., 2014), named *Tg(bactin:mCherry-dram1).* We could readily trace mCherry-Dram1 over time by confocal imaging epithelial cells in the thin tissue of the tail fin of 3 days post fertilisation (dpf) zebrafish larvae (Figure 1B and C). Time-lapse imaging revealed that mCherry-Dram1 labels motile and morphologically diverse globular and tubular vesicles (Figure 1C, Supplementary movie 1). We used a LysoTracker probe that accumulates and fluoresces in endolysosomal compartments with low luminal pH to confirm that mCherry-Dram1 mainly localises to these acidic organelles (Figure 1D). Previously we have demonstrated that ectopic activation of Dram1 by means of mRNA overexpression increased the number of autophagosomes observed per cell, and that transiently expressed mCherry-Dram1 can interact with autophagosomes (van der Vaart et al., 2014). To confirm that the mCherry-Dram1 construct retains the function of the endogenous protein, we crossed heterozygous *Tg(bactin:mCherry-dram1)* animals with the autophagy reporter line *Tg(CMV:GFP-Lc3)* (He et al., 2009). Co-expression of these two transgenes confirmed that mCherry-Dram1 interacts with autophagosomes and that ectopic expression of mCherry-Dram1 increased the number of autophagosomes observed per cell (Figure S1A and B). We can therefore use artificial expression of mCherry-Dram1 as a gain-of-function approach to study the role of Dram1 in vesicle trafficking.

To visualise endosomal vesicles that Dram1 interacts with, we used a transgenic line that fluorescently reports early endosomes in basal cell layer epithelial cells of the zebrafish epidermis: *TgBAC(ΔNp63:Gal4FF)^la213^; Tg(4xUAS:EGFP-2xFYVE)^la214^,* hereafter referred to as GFP-2xFYVE (Rasmussen et al., 2015). The GFP-2xFYVE probe incorporates specifically in membranes containing PI(3)P via its FYVE domains, thereby labelling early endosomes. However, a specific pool of PI(3)P also labels (nascent) autophagosomes (Nascimbeni et al., 2017). We therefore first tested the specificity of the GFP-2xFYVE probe by combining it with a *Tg(bactin:mCherry-Lc3)* line that marks autophagosomes, hereafter referred to as mCherry-Lc3. We found that GFP-2xFYVE and mCherry-Lc3 labelled autophagosomes rarely colocalise, but label distinct vesicles that occasionally are found in close proximity of each other (Figure S1C). Since the GFP-2xFYVE probe does not label autophagosomes, we therefore refer to vesicles labelled by GFP-2xFYVE in their membrane as ‘early endosomes’.

Confocal imaging of the GFP-2xFYVE and mCherry-Dram1 transgenes in the accessible zebrafish tail fin tissue allowed us to study endosomal dynamics in great detail. Time-lapse imaging demonstrated that globular mCherry-Dram1 labelled vesicles frequently interact with the PI(3)P-containing membrane of early endosomes (Figure 1E). We could also observe mCherry-Dram1 vesicles forming tethers between two distant early endosomes that are subsequently brought together (Figure 1E). Ultimately, mCherry-Dram1 labelled vesicles fuse with early endosomes and localise to their lumen. Early endosomes that have undergone such fusion events gradually lose the GFP-2xFYVE labelling of their membrane, representing a reduction of PI(3)P lipids present in these membranes. Taken together, ectopically expressed mCherry-Dram1 labels acidic and morphologically diverse vesicles that interact and fuse with early endosomes. Subsequently, these early endosomes alter the PI lipid composition of their membrane.

### Inhibiting PIKfyve and PI(3,5)P_2_ formation affects mCherry-Dram1 labelled vesicles

Early endosomes that have fused with mCherry-Dram1 labelled vesicles lose the GFP-2xFYVE labelling of their membrane. We hypothesised that the enzymatic activity of the 1-phosphatidylinositol 3-phosphate 5-kinase PIKfyve was responsible for the conversion of PI(3)P into PI(3,5)P_2_ in this process. To test this, we used YM201636 and apilimod to selectively inhibit the kinase activity of PIKfyve (Cai et al., 2013; Jefferies et al., 2008). A block in PIKfyve activity is known to affect fusion and fission events, which leads to membrane conservation and subsequent enlargement of endosomal compartments (Choy et al., 2018; Ikonomov et al., 2003; Sbrissa et al., 1999). As expected, GFP-2xFYVE zebrafish larvae treated with either YM201636 or apilimod displayed enlarged early endosomal vesicles marked by PI(3)P in their membranes (Figure S2A). As the more potent and selective of the two inhibitors (Cai et al., 2013), we tested a range of treatment durations for apilimod and found that a relatively short incubation of 2 hours robustly enlarged GFP-2xFYVE labelled vesicles (Figure S2B). We selected this treatment window for further experiments in which we exposed zebrafish larvae expressing both the GFP-2xFYVE and mCherry-Dram1 constructs to either apilimod or DMSO as a solvent control. We used confocal microscopy to image epithelial cells in the zebrafish tail fin and analysed the number and morphology of GFP-2xFYVE and mCherry-Dram1 vesicles per cell (Figure S3). Inhibition of the enzymatic activity of PIKfyve resulted in enlarged early endosomes and mCherry-Dram1 labelled vesicles, while the number of both types of vesicles per cell was reduced (Figure 2A-C). Furthermore, apilimod treatment significantly decreased the number of tubular mCherry-Dram1 vesicles per cell (Figure 2D & E). Therefore, mCherry-Dram1 labelled vesicles - that can interact and fuse with early endosomes - are reduced in number and altered in their morphology when PIKfyve is inhibited from converting PI(3)P on endosomal membranes into PI(3,5)P_2_.

**Figure 2:**
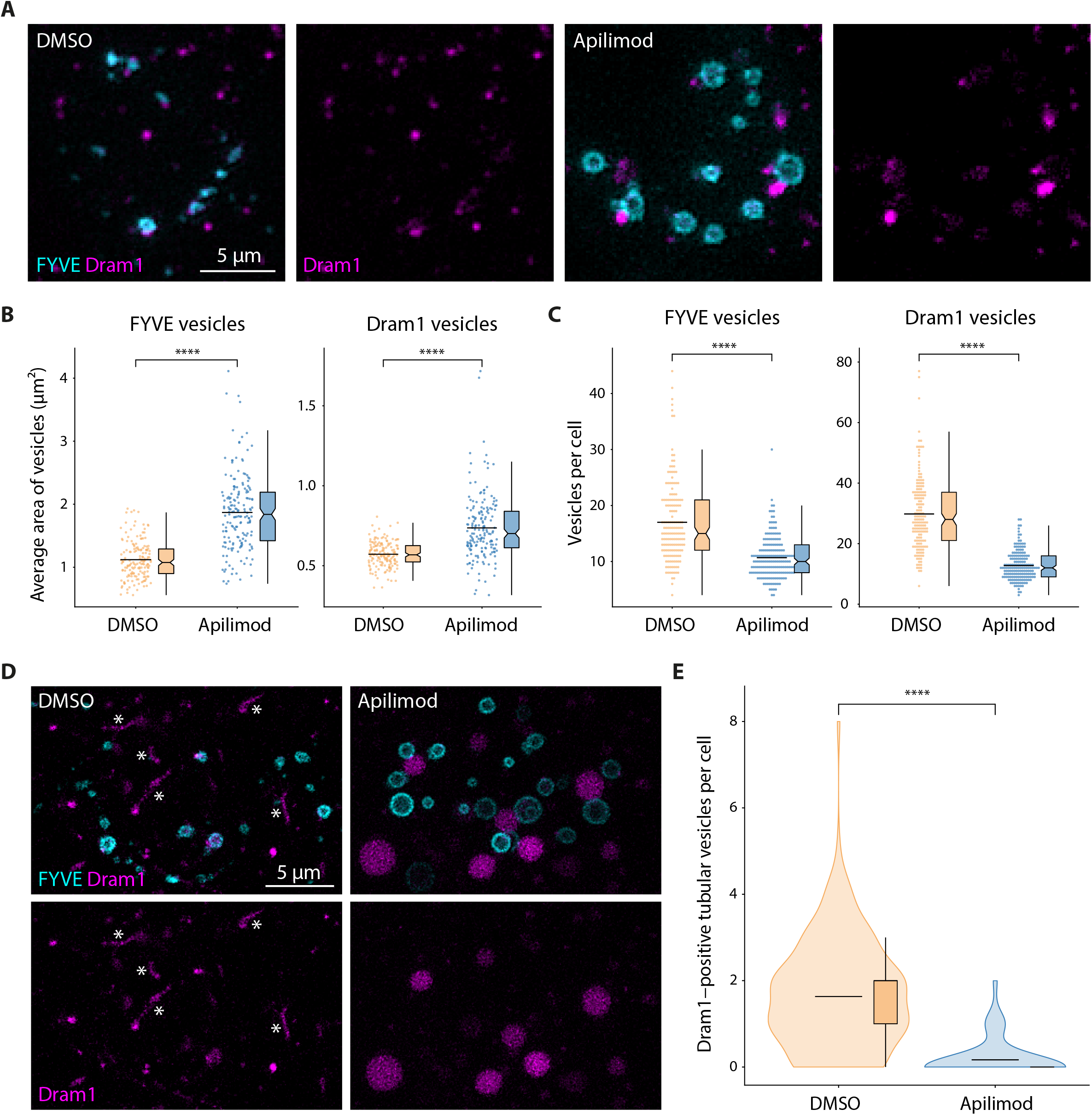
Inhibiting the formation of PI(3,5)P_2_ affects the morphology and number of mCherry-Dram1 labelled vesicles. Zebrafish larvae (3 dpf) expressing mCherry-Dram1 and GFP-2xFYVE were treated for 2 hours with 5 μm apilimod or DMSO as a solvent control. (A) Representative maximum intensity Z-projection of mCherry-Dram1 and GFP-2xFYVE in basal cell layer epithelial cells. The left hand panels display the merged image with mCherry-Dram1 in magenta and GFP-2xFYVE in cyan, while the right hand panels show only mCherry-Dram1. (B) Quantification of the average size of GFP-2xFYVE (FYVE) and mCherry-Dram1 (Dram1) labelled vesicles per basal cell layer epithelial cell. (C) Quantification of the number of GFP-2xFYVE (FYVE) and mCherry-Dram1 (Dram1) labelled vesicles per basal cell layer epithelial cell. (D) Maximum intensity Z-projection of mCherry-Dram1 and GFP-2xFYVE in basal cell layer epithelial cells displaying multiple tubular mCherry-Dram1 labelled vesicles (indicated by asterisks, *). (E) Quantification of the number of tubular mCherry-Dram1 labelled vesicles per basal cell layer epithelial cell. Tubular vesicles are defined as vesicles with a length that is at least two times longer than their width. Quantifications (B, C, E) were performed on n = 160 cells for the DMSO group and n = 173 cells for the apilimod treated group. For both conditions, these cells were imaged in the tailfins of 17 zebrafish larvae derived from 2 independent experiments. Scale bars: 5 μm.

### mCherry-Dram1 accumulates in the lumen and on the membrane of early endosomes upon inhibition of PI(3,5)P_2_ formation

Inhibiting PIKfyve to prevent the conversion of PI(3)P into PI(3,5)P_2_ altered the number and morphology of mCherry-Dram1 labelled vesicles, even though these vesicles themselves are initially devoid of PI(3)P. This observation suggests that the conversion of PI(3)P into PI(3,5)P_2_ on endosomal membranes is important for the fate or function of mCherry-Dram1 labelled vesicles. Therefore, we explored how inhibition of PIKfyve affects the localisation of mCherry-Dram1 and its interaction with early endosomes containing PI(3)P in their membranes. Based on confocal images of GFP-2xFYVE and mCherry-Dram1 in epithelial cells in the zebrafish tail fin, we could categorise mCherry-Dram1 signal into four groups: 1) mCherry-Dram1 signal that is distant from early endosomes; 2) mCherry-Dram1 signal that is in close proximity or directly adjacent to early endosomes; 3) mCherry-Dram1 signal that overlaps with the membrane of early endosomes; and 4) mCherry-Dram1 signal that is contained within early endosomes (Figure 3A). We then used Fiji/ImageJ to analyse the localisation of mCherry-Dram1 in respect to early endosomes according to these four categories (Figure S3). We found that PIKfyve inhibition reduced the number of times mCherry-Dram1 was localised distant from, adjacent to, or overlapping with early endosomal membranes, while it increased the number of times that mCherry-Dram1 was contained within early endosomes (Figure 3B). However, since inhibition of PIKfyve reduced the total number of mCherry-Dram1 vesicles per cell (Figure 2C), we also analysed the categories as a percentage of the total mCherry-Dram1 labelled vesicles present in each cell. We found that PIKfyve inhibition increased the percentage of mCherry-Dram1 signal that is contained within early endosomes or overlaps with early endosomal membranes, at the expense of the percentage of mCherry-Dram1 signal that is localised distant from or adjacent to early endosomes (Figure 3C). In conclusion, mCherry-Dram1 accumulates in the lumen of early endosomes and on their membranes when the conversion of PI(3)P into PI(3,5)P_2_ is inhibited.

**Figure 3:**
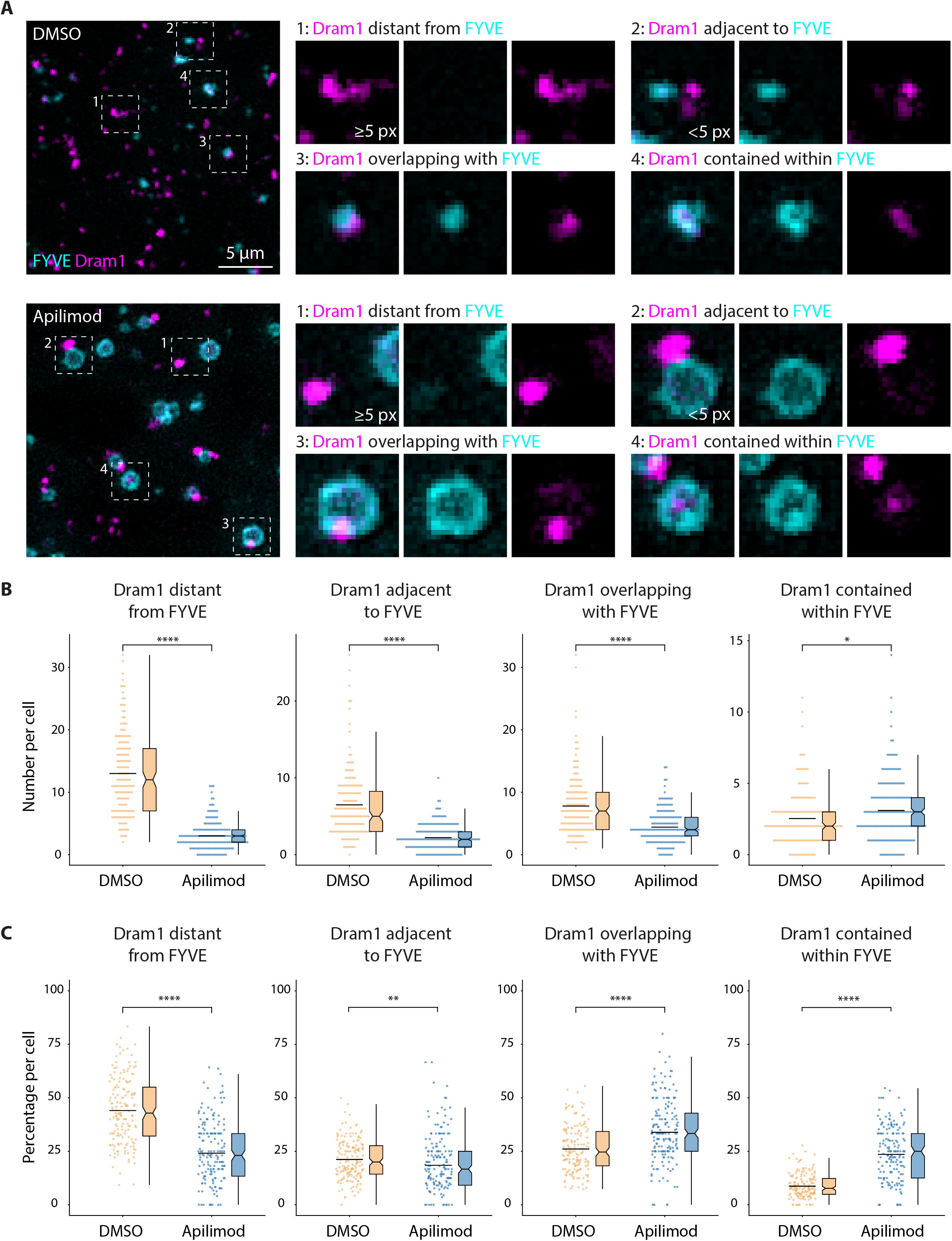
Dram1 accumulates in early endosomes and on early endosomal membranes upon inhibition of PI(3,5)P_2_ formation. Zebrafish larvae (3 dpf) expressing mCherry-Dram1 and GFP-2xFYVE were treated for 2 hours with 5 μm apilimod or DMSO as a solvent control. (A) Maximum intensity Z-projection of mCherry-Dram1 and GFP-2xFYVE in basal cell layer epithelial cells illustrating the 4 types of interactions that were categorised. Top panels: DMSO treated controls. Bottom panels: apilimod treated. Boxed areas in the merged images on the left hand side (numbered 1 to 4) are detailed on the right hand side, with mCherry-Dram1 in magenta and GFP-2xFYVE in cyan. (B) Quantification of the 4 types of interactions between mCherry-Dram1 and GFP-2xFYVE labelled vesicles per basal cell layer epithelial cell. (C) Quantification of the 4 types of interaction between mCherry-Dram1 and GFP-2xFYVE labelled vesicles per basal cell layer epithelial cell, displayed as percentage of the total number of mCherry-Dram1 labelled vesicles in a cell. Quantifications were performed on n = 160 cells for the DMSO group and n = 173 cells for the apilimod treated group. For both conditions, these cells were imaged in the tailfins of 17 zebrafish larvae derived from 2 independent experiments. Scale bars: 5 μm.

### Inhibition of PI(3,5)P_2_ formation reduces the dynamic interactions between mCherry-Dram1 and early endosomes

The observation that mCherry-Dram1 labelled vesicles accumulated in and on early endosomes upon inhibition of PIKfyve means that the dynamic interaction between these two types of vesicles was altered. Either mCherry-Dram1 labelled vesicles interacted more frequently with early endosomes upon inhibition of PIKfyve, or subsequent processes were inhibited that caused their accumulation. We performed time-lapse imaging of the interactions between mCherry-Dram1 and early endosomes to study these possible explanations. We exposed zebrafish larvae expressing both the GFP-2xFYVE and mCherry-Dram1 constructs to either apilimod or DMSO as a solvent control and imaged epithelial cells in the zebrafish tailfin after two hours of drug treatment using confocal microscopy (Figure 4A). In the control group, we observed many interactions between mCherry-Dram1 and early endosomes over time. This included temporary ‘kiss-and-run’ interactions, as well as long term contact between two or more vesicles which frequently ended in mCherry-Dram1 fusing into early endosomes (Supplementary movie 2). In contrast, inhibition of PIKfyve greatly reduced the motility of both types of vesicles, with interactions taking place infrequently and novel fusion events between mCherry-Dram1 and early endosomes only occurring rarely (Supplementary movie 3). Analysis of the number of interactions that took place with mCherry-Dram1 per early endosome confirmed our observations, as these were significantly reduced upon inhibition of PIKfyve (Figure 4B).

**Figure 4:**
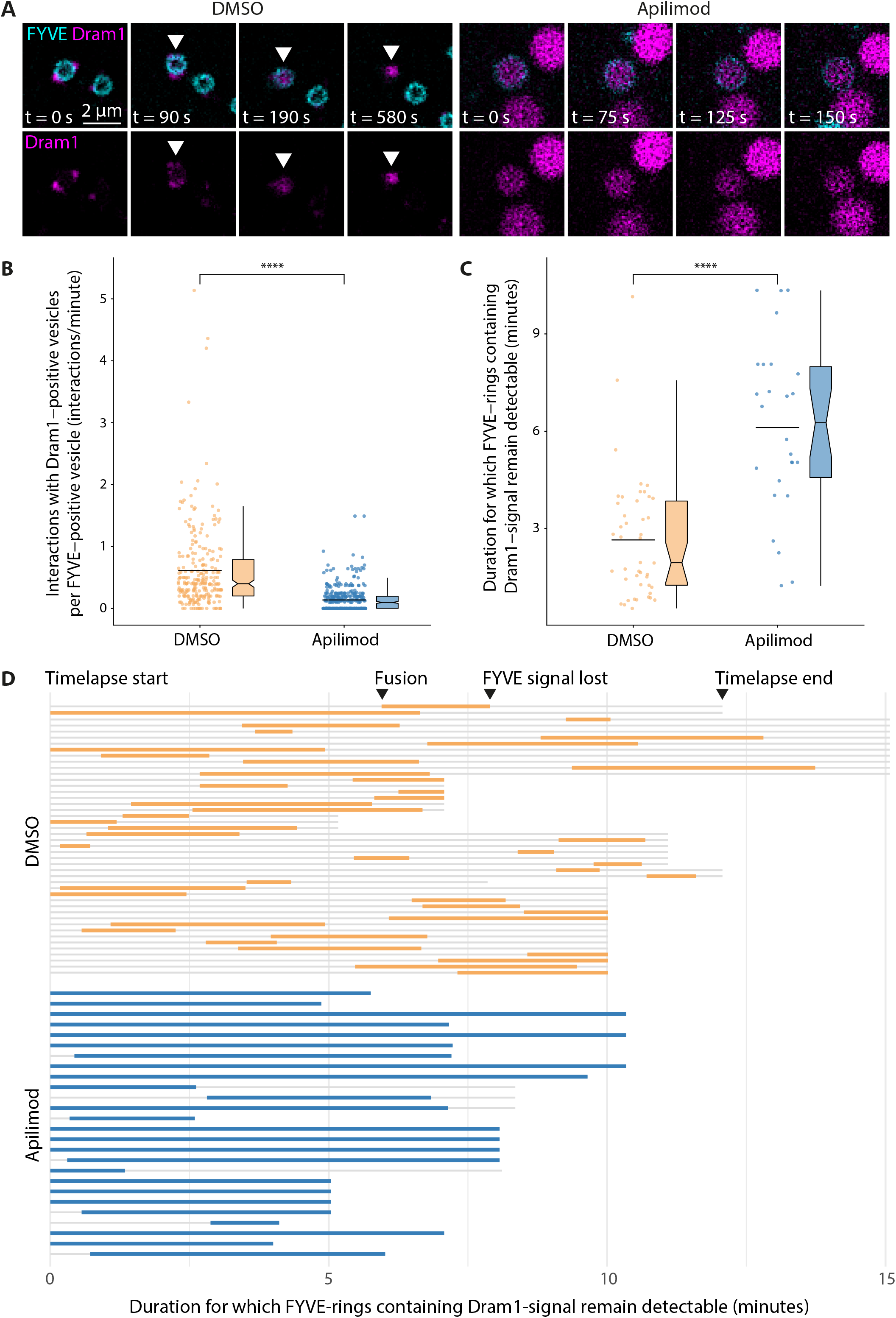
Interaction and fusion between Dram1-containing vesicles and early endosomes is reduced upon inhibition of PI(3,5)P_2_ formation. Zebrafish larvae (3 dpf) expressing mCherry-Dram1 and GFP-2xFYVE were treated for 2 hours with 5 μm apilimod or DMSO as a solvent control. (A) Representative stills from time-lapse confocal imaging of mCherry-Dram1 and GFP-2xFYVE in basal cell layer epithelial cells. The top panels display the merged image for each time point with mCherry-Dram1 in magenta and GFP-2xFYVE in cyan, while the bottom panels show only mCherry-Dram1. The arrowheads (Δ) indicate a GFP-2xFYVE labelled vesicle that fuses with mCherry-Dram1 (t=90s and t=190s) and subsequently loses the GFP-2xFYVE labelling of its membrane (t=580s). (B) Quantification of the number of observed interactions between mCherry-Dram1 and GFP-2xFYVE labelled vesicles per minute. For the DMSO control group, the interactions of n = 248 GFP-2xFYVE labelled vesicles with mCherry-Dram1 imaged in 29 different cells were quantified for the duration of the time lapses. For the apilimod treated group, the interactions of n = 341 GFP-2xFYVE labelled vesicles with mCherry-Dram1 imaged in 40 different cells were quantified for the duration of the time lapses. (C) Quantification of the duration for which GFP-2xFYVE labelling of membranes could be observed following fusion with mCherry-Dram1 labelled vesicles (DMSO: n = 45 fusion events in 29 cells; apilimod: n = 26 fusion events in 40 cells). (D) Visualisation of the duration for which GFP-2xFYVE labelling of membranes could be observed following fusion with mCherry-Dram1 labelled vesicles, relative to the length of time for which the vesicle could be imaged. Horizontal light-grey bars indicate the length of time for which the cell could be imaged. A yellow (DMSO) or blue (apilimod) horizontal bar indicates the moment of fusion, up until the moment the GFP-2xFYVE labelling of the membrane could no longer be observed. Scale bars: 5 μm.

The anticipated effect of PIKfyve inhibition is that PI(3)P in the membrane of early endosomes can no longer be converted into PI(3,5)P_2_. As observed before (Figure 1E), early endosomes in the control group gradually lost the PI(3)P lipids marked by GFP-2xFYVE in their membrane following fusion events with mCherry-Dram1 (Figure 4A, Supplementary movie 2). Upon inhibition of PIKfyve, early endosomes that had already fused with mCherry-Dram1, or underwent novel fusion events on rare occasions, no longer lost the GFP-2xFYVE labelling of their membranes (Supplementary movie 3). By quantifying this process for multiple time-lapse recordings, we confirmed that the duration for which GFP-2xFYVE labelling of early endosomal membranes remained detectable following fusion with mCherry-Dram1 vesicles was significantly increased upon PIKfyve inhibition (Figure 4C). Not all time-lapse recordings of epithelial cells in zebrafish tail fins were of equal length due to technical difficulties associated with this type of imaging in live animals (e.g. samples drifting out of focus). We therefore also plotted the duration for which a GFP-2xFYVE ring containing mCherry-Dram1 signal remained detectable in relation to the total duration for which the cell in which the fusion event occurred could be followed (Figure 4D). This visualisation clearly illustrates the difference between the control group in which mCherry-Dram1 frequently fused with early endosomes that subsequently lost the GFP-2xFYVE labelling of their membrane, and the apilimod treated group in which the majority of early endosomes containing mCherry-Dram1 signal retain the GFP-2xFYVE labelling of their membrane for the entire duration of the time-lapse. Taken together, inhibition of PI(3,5)P_2_ formation reduced the dynamic interactions between mCherry-Dram1 and early endosomes, and caused mCherry-Dram1 to accumulate in early endosomes by halting processes that normally follow upon vesicle fusion.

### Acidification of mCherry-Dram1 vesicles is reduced upon inhibition of PI(3,5)P_2_ formation

We have thus shown that early endosomes that have fused with mCherry-Dram1 labelled vesicles lose the GFP-2xFYVE labelling of their membrane in a process dependent on the kinase activity of PIKfyve. This loss of signal suggests that an endosomal maturation process takes place in which PI(3)P is converted into PI(3,5)P_2_ present in late endosomal membranes. The maturation of early into late endosomes is associated with a decrease in luminal pH (Maxfield and Yamashiro, 1987). This prompted us to investigate how inhibition of PIKfyve affected the acidification of early endosomes and mCherry-Dram1 labelled vesicles. We therefore imaged GFP-2xFYVE and mCherry-Dram1 in epithelial cells in the zebrafish tail fin, combined with LysoTracker staining to label acidic vesicles. In the control group, we observed that the majority of mCherry-Dram1 labelled vesicles are acidic (Figure 5A), confirming our earlier findings (Figure 1D and (van der Vaart et al., 2014)). GFP-2xFYVE labelled vesicles in the control group varied in the extent of their acidity, ranging from (almost) no detectable LysoTracker staining to clear staining of their lumen (Figure 5A). This variation in acidity for PI(3)P labelled vesicles likely reflects the gradual acidification of early endosomes that takes place as they mature. Upon inhibition of PIKfyve by apilimod treatment, early endosomes continue to display this range of luminal acidification, with smaller PI(3)P labelled vesicles frequently not or dimly stained by LysoTracker and larger vesicles typically stained intensely (Figure 5B). In contrast, mCherry-Dram1 labelled vesicles appeared to be less frequently and less intensely stained by LysoTracker when PIKfyve was inhibited (Figure 5B). We used Fiji/ImageJ to analyse the spatial overlap (colocalisation) between mCherry-Dram1 and LysoTracker staining and found that the correlation between these two fluorescent signals decreased significantly upon inhibition of PIKfyve (Figure 5C). We therefore conclude that the acidification of mCherry-Dram1 vesicles is at least partially dependent on the formation of PI(3,5)P_2_ by PIKfyve.

**Figure 5:**
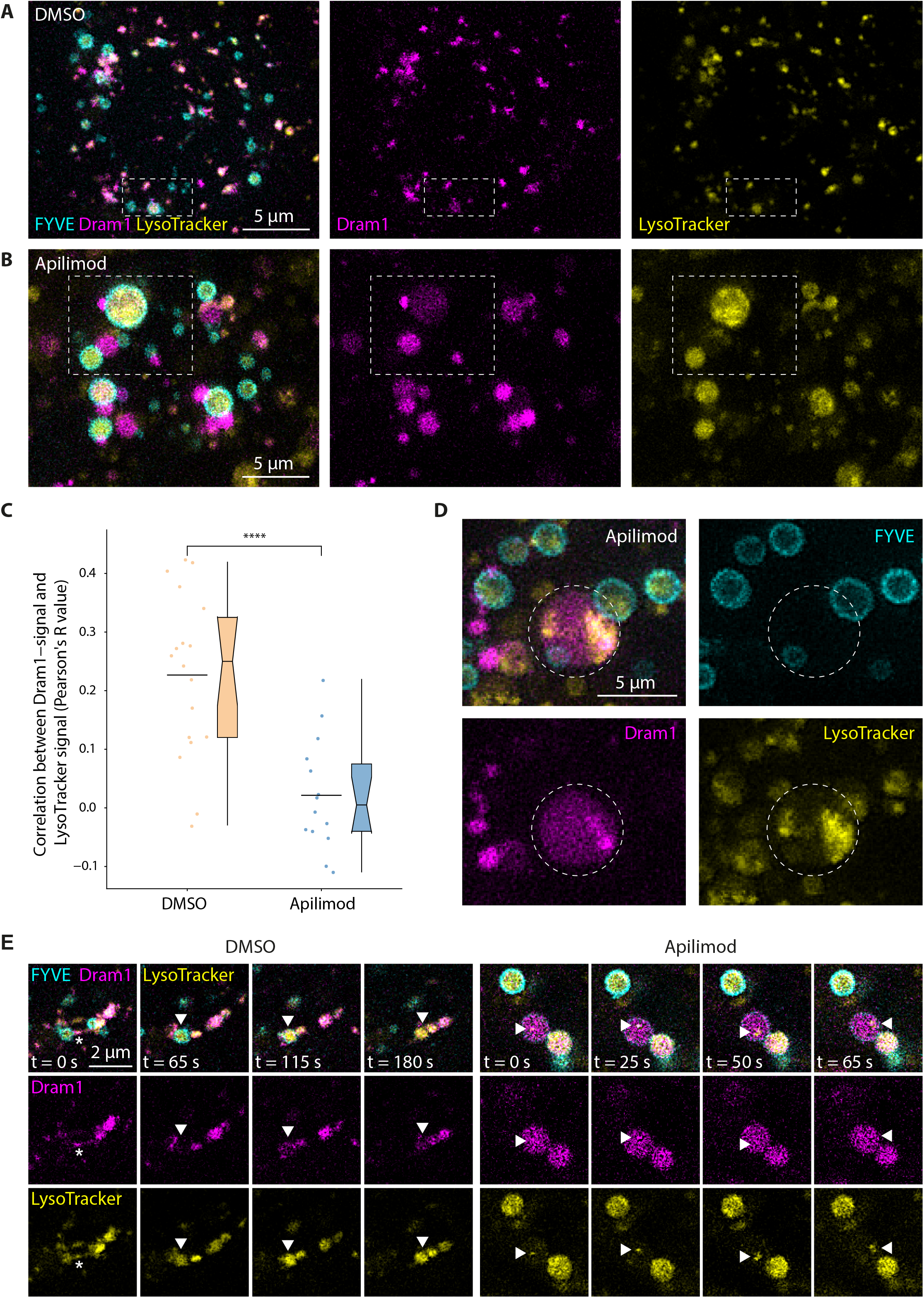
Acidification of Dram1-containing vesicles is reduced upon inhibition of PI(3,5)P_2_ formation, arresting Dram1 in early endosomes and MVBs. Zebrafish larvae (3 dpf) expressing mCherry-Dram1 and GFP-2xFYVE and stained with LysoTracker Deep Red were treated for 2 hours with 5 μm apilimod or DMSO as a solvent control. (A, B) Representative maximum intensity Z-projection of mCherry-Dram1, GFP-2xFYVE, and LysoTracker Deep Red in basal cell layer epithelial cells. The left hand panels display the merged image with mCherry-Dram1 in magenta, GFP-2xFYVE in cyan, and LysoTracker in yellow, while the middle and right hand panels show only mCherry-Dram1 and LysoTracker, respectively. The boxed area in the DMSO panels indicates GFP-2xFYVE labelled vesicles with LysoTracker staining ranging from dim to intense. The boxed area in the apilimod panels indicates mCherry-Dram1 labelled vesicles with LysoTracker staining ranging from dim to intense. (C) The Pearson’s R value correlation between mCherry-Dram1 and LysoTracker Deep Red fluorescent signal was determined for confocal images of basal cell layer epithelial cells in the tailfin of n = 18 (DMSO) and n = 14 (apilimod) zebrafish larvae derived from two independent experiments. Each of these images contained multiple epithelial cells. (D) Maximum intensity Z-projection of mCherry-Dram1, GFP-2xFYVE, and LysoTracker Deep Red in a basal cell layer epithelial cell treated with apilimod. The encircled area indicates a mCherry-Dram1 labelled compartment containing (remnants of) other vesicles positive for either GFP-2xFYVE or LysoTracker Deep Red. (E) Representative stills from time-lapse confocal imaging of mCherry-Dram1, GFP-2xFYVE, and LysoTracker Deep Red in basal cell layer epithelial cells. The top panels display the merged image for each time point with mCherry-Dram1 in magenta, GFP-2xFYVE in cyan, and LysoTracker Deep Red in yellow. The middle and bottom panels show only mCherry-Dram1, or LysoTracker Deep Red respectively. A tether formed by mCherry-Dram1 between two GFP-2xFYVE labelled vesicles is indicated by an asterix (*). The arrowheads (Δ) in DMSO panels indicate a GFP-2xFYVE labelled vesicle that fuses with mCherry-Dram1 (t=65s and t=115s) and subsequently loses the GFP-2xFYVE labelling of its membrane while increasing the intensity of its LysoTracker Deep Red staining (t=180s). The arrowheads (Δ) in apilimod panels indicate a LysoTracker Deep Red stained intraluminal vesicle moving inside a mCherry-Dram1 labelled compartment. Scale bars: 5 μm (A, B and D) or 2 μm (E).

While analysing the colocalisation between mCherry-Dram1 and LysoTracker, we encountered multiple large mCherry-Dram1 labelled vesicles that contained acidic (Lysotracker stained) and non-acidic GFP-2xFYVE labelled vesicles (Figure 5D). These intraluminal vesicles appeared to accumulate within the mCherry-Dram1 labelled compartments, forming what resembles a multivesicular body. To visualise the dynamics of these events, we performed time-lapse imaging of GFP-2xFYVE and mCherry-Dram1 combined with LysoTracker staining. In the control situation, a mCherry-Dram1^+^/LysoTracker^+^ vesicle formed a tether between two early endosomes with dim LysoTracker staining, causing the two early endosomes to fuse together (Figure 5E and Supplementary movie 4). The mCherry-Dram1^+^/LysoTracker^+^ vesicle continued to interact with this newly formed endosome and ultimately fused with it. Following this fusion event, the early endosome displayed more intense luminal LysoTracker staining and lost the GFP-2xFYVE labelling of its membrane over time. This maturation process forms a stark contrast to what occurred upon inhibition of PIKfyve. As described before (Figure 4A), mCherry-Dram1 and GFP-2xFYVE labelled vesicles rarely interacted nor altered their existing associations (Figure 5E and Supplementary movie 5). Large mCherry-Dram1 labelled vesicles varied in their acidity, ranging from no or dim LysoTracker staining to intense LysoTracker staining. Inside the lumen of non-acidified mCherry-Dram1 labelled compartments, we regularly observed small acidic vesicles that moved around in a seemingly random pattern (Figure 5E and Supplementary movie 5). These acidic intraluminal vesicles persisted over time, with no indication of releasing their content into the lumen in which they reside. In conclusion, the kinase activity of PIKfyve is required for mCherry-Dram1 labelled vesicles to tether early endosomes and fuse with them to kickstart a maturation process in which their signature PI(3)P membrane lipids are converted and their lumen further acidifies. When PIKfyve is inhibited, targeting of mCherry-Dram1 to acidic vesicles is reduced, arresting mCherry-Dram1 in multivesicular bodies, early endosomes, or non-acidified vesicles halted in their fusion with early endosomes.

## Discussion

Degrading unwanted or harmful elements present in a cell or its microenvironment is important to maintain cellular and tissue homeostasis. For instance, pathogenic protein aggregates can cause diseases like Alzheimer’s, Parkinson’s, or Huntington’s. Enhancing the delivery of pathogenic proteins to the degradative environment of the lysosome is one possible therapeutic approach for these protein aggregation diseases (Aguzzi and O’Connor, 2010). During microbial infection, degradation of pathogens in lysosomes is a key immune defence function and enhancing the underlying vesicle trafficking processes therefore presents a major opportunity for therapeutic strategies (Kaufmann et al., 2018). For both examples, a thorough understanding of the molecular mechanisms controlling endolysosomal and autophagic trafficking is required to successfully intervene in disease pathogenesis. Here, we add to our understanding of these processes by studying the function of Dram1 in vesicle trafficking in the optically transparent zebrafish model. We found that Dram1-mediated vesicle fusion is dependent on the formation of PI(3,5)P_2_ by PIKfyve to deliver vesicles and their cargo to the acidic environment of endolysosomes.

The interplay between Dram1 and PIKfyve revealed by our study sheds light on the molecular mechanisms underlying functions of Dram1 described in previous reports. Studies on mammalian cells and in the zebrafish model found that DRAM1 (Dram1 in zebrafish) can induce autophagy and stimulate vesicle fusion with lysosomes (Crighton et al., 2006; van der Vaart et al., 2014; Zhang et al., 2013). This role of Dram1 is important in defence against mycobacterial infection in the zebrafish model for tuberculosis, and mammalian DRAM1 was found to associate with *Mycobacterium tuberculosis* phagocytosed by primary human macrophages (van der Vaart et al., 2014). Furthermore, we recently described how zebrafish macrophages lacking Dram1 failed to deliver pathogenic mycobacteria to acidic endolysosomal compartments, ultimately resulting in an inflammatory type of cell death - called pyroptosis - which disseminates the infection (Zhang et al., 2020). Besides its function in vesicle trafficking, DRAM1 is required for apoptosis mediated by the tumour suppressor p53 in relation to cancer and in HIV infected CD4(+) T cells (Crighton et al., 2006; Laforge et al., 2013). DRAM1 was shown to interact with the pro-apoptotic protein BAX, which recruited BAX to lysosomes and initiated cell death via release of lysosomal cathepsin B (Guan et al., 2015). Recently, it has also been found that DRAM1 is required for efficient activation of mTORC1, a nutrient-sensing complex that functions at the lysosome (Beaumatin et al., 2019). DRAM1 facilitates activation of mTORC1 by binding the membrane carrier protein SCAMP3 and the amino acid transporters SLC1A5 and LAT1, thereby directing them to lysosomes (Beaumatin et al., 2019). An emerging theme is that DRAM1 functions at the interface between lysosomes, signalling complexes, and other vesicles by binding and directing effector molecules. Our *in silico* analysis of predicted protein domains further supports a role for DRAM1 as a protein binding hub important in the regulation of vesicle trafficking. Although Dram1-mediated vesicle fusion and maturation events required the enzymatic activity of PIKfyve to generate PI(3,5)P_2_ on endosomal membranes, it remains unclear whether Dram1 directly interacted with these molecules. Since the DRAM1 protein lacks consensus PI binding motifs (e.g. a FYVE domain), we expect that effector proteins capable of binding either PI(3)P or PI(3,5)P_2_ mediate this interaction.

The primary function of PIKfyve is to bind PI(3)P on endosomal membranes through its FYVE domain and phosphorylate it into PI(3,5)P_2_ (Shisheva, 2001). Besides this, PIKfyve can also phosphorylate PI to generate PI(5)P, a low abundant PI family member found in different cellular compartments, including the nucleus (Poli et al., 2019; Shisheva, 2001). PIKfyve functions as part of a complex scaffolded by VAC14, also known as ArPIKfyve (Associated Regulator of PIKfyve) (Sbrissa et al., 2004). This complex also contains Sac3, the phosphatase that converts PI(3,5)P_2_ into PI(3)P (Sbrissa et al., 2008). The presence of two enzymes with opposing activities in the same complex indicates that PI(3,5)P_2_ levels need to be tightly controlled. Indeed, inactivation of the PIKfyve containing complex impaired autophagic and endolysosomal vesicle trafficking, thereby halting the maturation of these vesicles (de Lartigue et al., 2009; Dong et al., 2010; Ferguson et al., 2009; Kim et al., 2014). The typical enlargement of lysosomes upon inhibition of PIKfyve is ascribed to lysosome coalescence, most likely due to reduced fission events during lysosomal ‘kiss-and-run’ interactions and/or full fusion and fission cycles (Choy et al., 2018). Based on the data presented here, we propose that inhibition of PIKfyve prevents DRAM1 from performing its function as an interface between lysosomes and vesicles destined for fusion with lysosomes.

We took advantage of a fluorescently tagged version of zebrafish Dram1 to study its dynamic localisation during vesicle trafficking events. This approach yielded valuable insights into the role of DRAM1 in the endolysosomal maturation process, demonstrating that early endosomes labelled by PI(3)P in their membrane mature and acidify following fusion events with mCherry-Dram1 labelled vesicles. The overexpression of mCherry-Dram1 (driven by the zebrafish beta actin promoter) mimics situations in which cells have upregulated the expression of DRAM1 in response to cellular stressors like DNA damage or infection. However, this approach comes with a number of caveats regarding the interpretation of our results, since ectopically expressed tagged proteins can exhibit altered behaviour or expression patterns compared to their endogenous counterparts. For this reason, we place less emphasis on the identity of mCherry-Dram1 labelled vesicles and rather focus on their activity and interactions. Faithfull determination of the subcellular localisation of DRAM1 under different circumstances would require an antibody staining approach to detect the endogenous protein, which would preclude any dynamic observations. Furthermore, the expression of fluorescently tagged proteins can alter cellular functions. It is known that expression of GFP-2xFYVE can alter endosomal dynamics and induce sustained autophagosome formation (Nascimbeni et al., 2017). Therefore we expect that GFP-2xFYVE interfered with PI(3)P interactions to some extent in our experiments, but the strong effect of PIKfyve inhibition on endosomal dynamics indicates that most of the PI(3)P functionality remains intact in the GFP-2xFYVE line. For mCherry-Dram1, we confirmed that the relatively large fluorescent tag did not interfere with its known localisation to acidic vesicles, nor its ability to induce autophagy upon overexpression. Nonetheless, a long sought-after goal in cell biology remains to study endogenous protein dynamics in live cells without altering their functionality, localisation, or expression level. Specifically for DRAM1, we aim to determine the function, identity, and dynamics of globular and tubular vesicles containing endogenous DRAM1 in their membrane or lumen.

Based on our observations, we conclude that mCherry-Dram labelled vesicles can tether early endosomes and fuse with them as part of their maturation process. When we inhibited the formation of PI(3,5)P_2_ by PIKfyve, targeting of mCherry-Dram1 to acidic endolysosomal vesicles was reduced, strongly suggesting that cargo carried by mCherry-Dram1 labelled vesicles is destined for degradation in lysosomal compartments. In the zebrafish model for tuberculosis, we have previously demonstrated that overexpression of Dram1 enhanced the localisation of mycobacteria to acidic endolysosomes (van der Vaart et al., 2014). Further studies on the molecular mechanisms behind DRAM1-mediated vesicle trafficking events will hopefully help to understand how it targets cargo to the degradative environment of the lysosome. Such knowledge could form the basis for therapeutic approaches for a spectrum of diseases in which unwanted elements reside inside a cell or in its microenvironment.

## Material & methods

### Zebrafish husbandry and care

Zebrafish lines in this study (listed in Supplementary table 1) were handled in compliance with local animal welfare regulations, as overseen by the Animal Welfare Body of Leiden University (License number: 10612) and maintained according to standard protocols (zfin.org). All experiments were performed on embryos or larvae up to 3 days post-fertilization (dpf), which have not yet reached the free-feeding stage. Embryos/larvae were kept in egg water (60 μg/ml Instant Ocean sea salts) at 28.5°C and treated with 0.02% ethyl 3-aminobenzoate methanesulfonate (Tricaine, Sigma-Aldrich) for anesthesia before imaging and fixation. For all experiments involving *Tg(bactin:mCherry-dram1)*, female adult zebrafish heterozygous for the transgene were outcrossed with male adult zebrafish of the required genotype (e.g. AB/TL wild type or carrying the GFP-x2FYVE transgenic construct). Offspring of these crosses were selected for proper expression of the transgenic constructs at 2 dpf by stereo fluorescent microscopy.

### Generation of transgenic reporter lines

Full-length zebrafish Lc3 cDNA *(map1lc3b-201;* ENSDART00000163508.2) with attB sites added to its sequence was synthesised (BaseClear) and used to create a 3’ Gateway entry vector (Invitrogen). This 3’ Gateway entry vector was combined into a Tol2 containing destination vector together with a 5’ Gateway entry vector containing the zebrafish beta actin promoter and a Gateway middle entry vector containing mCherry with the stop codon removed, generating the following DNA construct: *bactin:mCherry-Lc3.* For the generation of *Tg(bactin:mCherry-dram1),* we used a DNA construct that was previously generated (van der Vaart et al., 2014). The DNA constructs were injected into AB/TL wildtype zebrafish embryos at the one cell stage (1 nl at 50 ng/μl), together with 50 pg Tol2 transposase mRNA to allow efficient integration into the genome. Zebrafish larvae were screened for appropriate expression of the constructs by stereo microscopy and reared into adulthood.

### Drug treatment

Larvae were bath treated with apilimod (S6414, Selleck) or YM201636 (S1219, Selleck) diluted into egg water at a working concentration of 5 μM or 10 μM, respectively.

### LysoTracker staining

Larvae were immersed in egg water containing 5 μM LysoTracker Deep Red (L12492, ThermoFisher) for 1 hour. Embryos were rinsed 3 times with egg water before imaging.

### Confocal laser scanning microscopy

When appropriate, larvae were fixed in 4% formaldehyde (28906, ThermoFisher) in PBS solution overnight at 4°C. Fixed or Live embryos were mounted with 1.5% low melting agarose (140727, SERVA) in PBS or egg water, respectively. Basal cell layer epithelial cells were imaged in the thin and optically transparent tail fin area using a Leica TCS SP8 confocal microscope with a 63X oil immersion objective (NA = 1.4), and equipped with 488 nm, 532 nm, and 638 nm laser lines. For time-lapse imaging, confocal micrographs were acquired for a single focal plane at a time interval of ~1.3 second/image. Representative images were deconvoluted using the Iterative Deconvolution 3D plugin in Fiji/ImageJ (Dougherty, 2005).

### Image analysis

Raw imaging data was analysed in Fiji/ImageJ to obtain measurements for vesicle morphology, interactions between vesicles, and colocalisation of fluorescent signals. For measurements of vesicle morphology, a maximum intensity Z-projection was generated for a single layer of epithelial cells imaged in the zebrafish tailfin tissue. Individual cells were selected and stored as regions of interest (ROIs) using the Polygon selection tool. The Phansalkar Auto-Local Threshold method was used for segmentation of vesicles. Segmented vesicles that were directly adjacent to each other were separated using a Watershed function. The resulting individual vesicles were measured per cell using the Analyze Particles function.

To measure interactions between vesicles, the same method as described above was used to segment individual vesicles per cell. Vesicles labelled by their respective fluorescent signal were stored as ROIs. Subsequently, the distance between each mCherry-Dram1 ROI and the nearest GFP-2xFYVE labelled ROI was determined. Based on this measurement, mCherry-Dram1 ROIs were categorised into four groups: 1) mCherry-Dram1 ROI that is distant from a GFP-2xFYVE ROI (distance ≥ 5 pixels); 2) mCherry-Dram1 ROI that is in close proximity or directly adjacent to a GFP-2xFYVE ROI (distance < 5 pixels); 3) mCherry-Dram1 ROI that overlaps with a GFP-2xFYVE ROI; and 4) mCherry-Dram1 ROI that is contained within a GFP-2xFYVE ROI. The Fiji/ImageJ plugin created to automate this analysis, called ‘FYVE DRAM Analysis’, is openly available for download via the Leiden University update site (http://sites.imagej.net/Willemseji/).

To analyse colocalisation between mCherry-Dram1 and LysoTracker Deep Red fluorescent signals, a maximum intensity Z-projection was generated for a single layer of epithelial cells imaged in the zebrafish tailfin tissue. The Gaussian Blur (sigma = 1) function was applied to decrease noise. After this, the Li Threshold method, followed by the Analyze Particles function (‘Show Mask’; size cut off of 10 pixels) was used to create a binary mask that excludes zero-zero pixels from the colocalisation analysis. Finally, we used the Coloc 2 Fiji/ImageJ plugin (available via https://imagej.net/Coloc_2) to determine the Pearson correlation coefficient between the two fluorescent signals.

### Statistical analysis and data representation

Statistical analyses were performed using GraphPad Prism software (Version 5.01; GraphPad). All experimental data (mean ± SEM) was analyzed using unpaired, two-tailed Mann-Whitney U tests for comparisons between two groups and Kruskal-Wallis one-way analysis of variance with Dunn’s multiple comparison methods as a posthoc test for comparisons between more than two groups. (ns, no significant difference; *p < 0.05; **p < 0.01; ***p < 0.001; ****p < 0.0001). The data sets from each group are shown in a scatter plot (left) and a boxplot (right). In the scatter plots each dot represents a data point, with the mean indicated by a horizontal line. Boxplots include 50% of the data points, with a vertical line indicating the 95% confidence interval and a horizontal line indicating the median. The only exception to this is Figure 2E, in which a violin plot is shown to represent the spread of individual data points due to the large number of 0 values which would distort the scatter plot.

## Acknowledgements

We thank Monica Varela and Rubén Marín Juez for critical proof reading of the manuscript. We are grateful to all members of the fish facility team for zebrafish care. M.v.d.V. was supported by the Netherlands Technology Foundation TTW (project 13259). S.M.S. was funded by the European Union’s Horizon 2020 research and innovation programme under the Marie Sklodowska-Curie grant agreement No. 721537. G.F.-C. was funded by a European Marie Curie fellowship (H2020-COFUND-2015-FP707404).

## Figure legends

**Supplementary figure 1.**
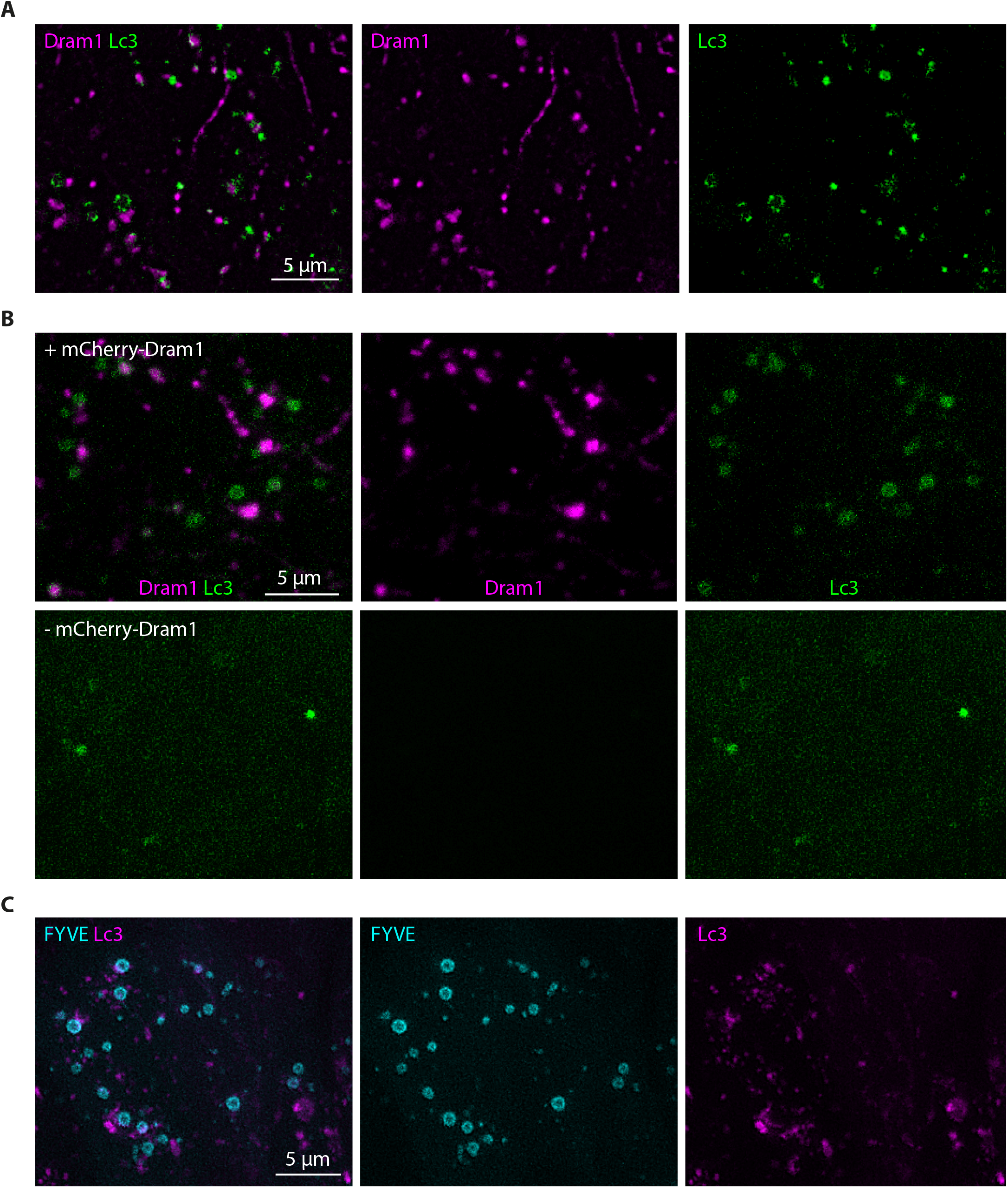
supporting Figure 1: Representative maximum intensity Z-projections of basal cell layer epithelial cells imaged in the tailfin of 3 days post fertilisation (dpf) zebrafish larvae. (A) Epithelial cells expressing mCherry-Dram1 and GFP-Lc3. Panels (from left to right) display the merged image, mCherry-Dram1 in magenta, and GFP-Lc3 in green. (B) Same as described for (A), with the exception that the offspring of heterozygous *Tg(bactin:mCherry-dram1)* animals outcrossed with *Tg(CMV:GFP-Lc3)* animals was sorted into groups that were either positive or negative for the mCherry-Dram1 construct, while all expressed the GFP-Lc3 construct. Top panels: expressing the mCherry-Dram1 construct (+ mCherry-Dram1). Bottom panels: not expressing the mCherry-Dram1 construct (-mCherry-Dram1). (C) Epithelial cells expressing mCherry-Lc3 and GFP-2xFYVE. Panels (from left to right) display the merged image, GFP-2xFYVE in cyan and mCherry-Lc3 in magenta. Scale bars: 5 μm.

**Supplementary figure 2.**
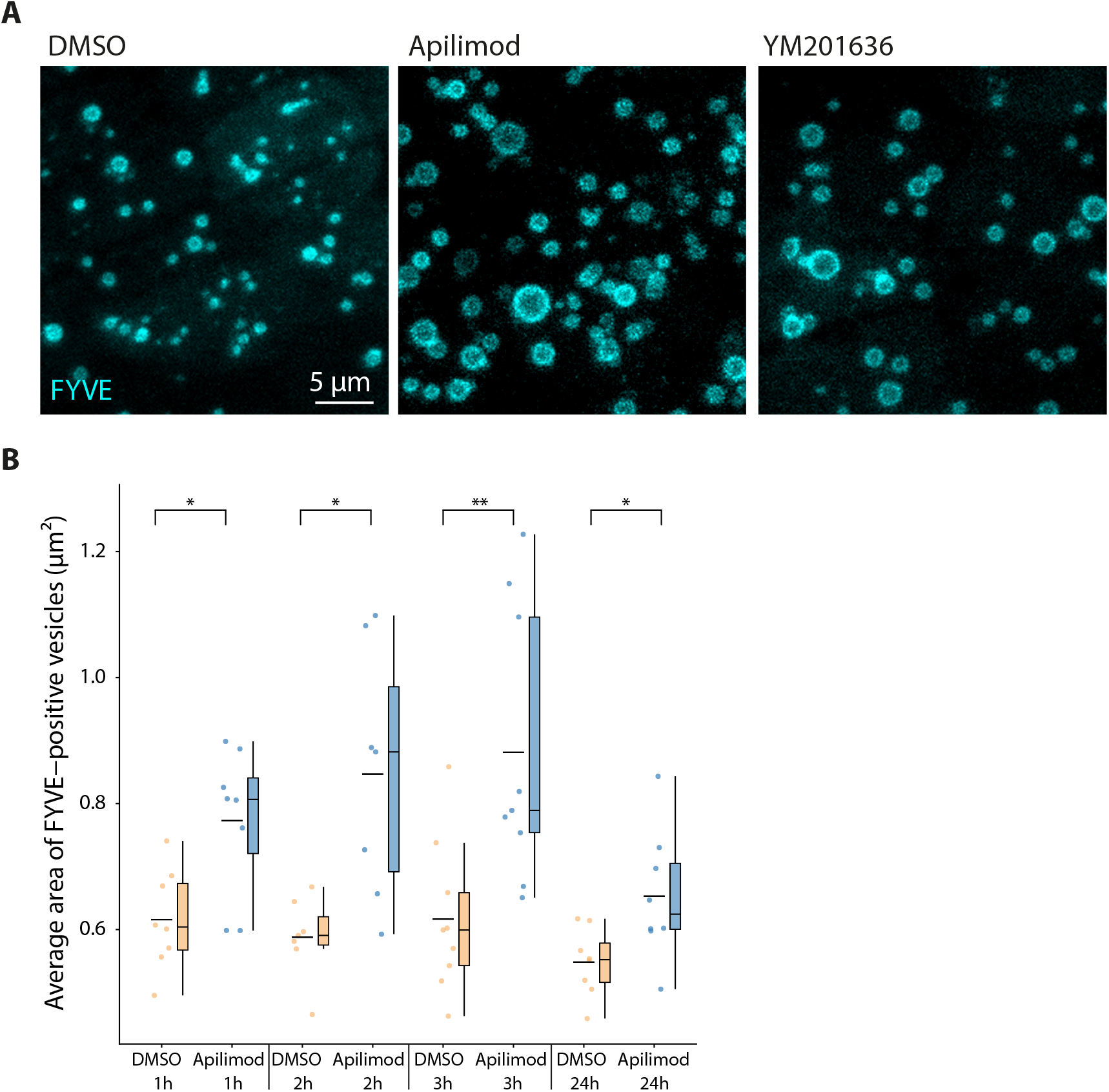
supporting Figure 2: (A) Zebrafish larvae (3 dpf) expressing GFP-2xFYVE were treated for 2 hours with 5 μm apilimod, 10 μm YM201636, or DMSO as a solvent control. Representative maximum intensity Z-projection of GFP-2xFYVE in basal cell layer epithelial cells. (B) Zebrafish larvae (3 dpf) expressing GFP-2xFYVE were treated for 1, 2, 3, or 24 hours prior to fixation and imaging with 5 μm apilimod or DMSO as a solvent control. The average area of GFP-2xFYVE labelled vesicles per cell was measured using Fiji/ImageJ. N ≥ 7 individual zebrafish larvae per group.

**Supplementary figure 3.**
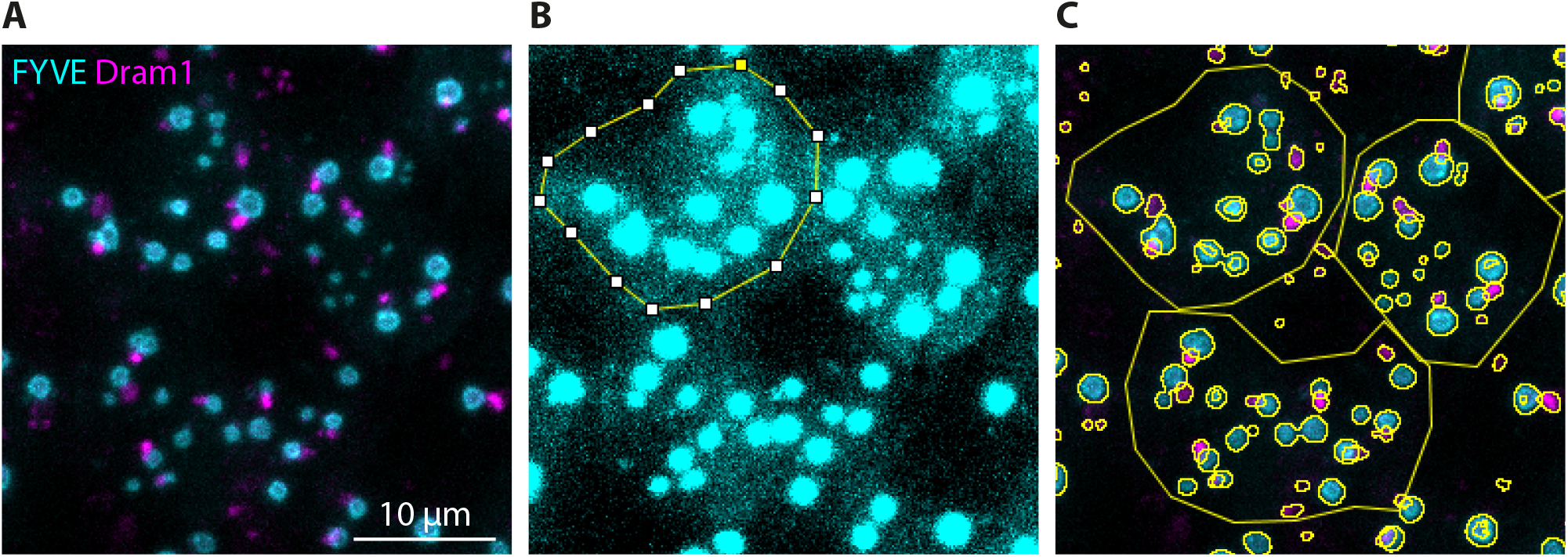
supporting Figure 2: (A) Representative maximum intensity Z-projection of basal cell layer epithelial cells expressing mCherry-Dram1 and GFP-2xFYVE, imaged in the tailfin of a 3 days post fertilisation (dpf) zebrafish larvae. (B) Example of a manually segmented epithelial cell based on a high-intensity representation of GFP-2xFYVE signal present in the cell. (C) Example of vesicle segmentation as performed by Fiji/Image.

**Supplementary table 1:** Zebrafish lines used in this study

| Line | Description | Reference |
| --- | --- | --- |
| AB/TL | Wild type strain | - |
| *TgBAC (ΔNp63:Gal4FF)<sup>la213</sup>* | Gal4 driver line specific for basal cell layer epithelial cells | (Rasmussen et al., 2015) |
| *Tg(4xUAS:EGFP-2xFYVE)<sup>la214</sup>* | Fluorescent probe labelling PI(3)P membrane lipids | (Rasmussen et al., 2015) |
| *Tg(CMV:GFP-Lc3)* | GFP-tagged zebrafish Lc3 | (He et al., 2009) |
| *Tg(bactin:mCherry-Lc3)* | mCherry-tagged zebrafish Lc3 | This study |
| *Tg(bactin:mCherry-dram1)* | mCherry-tagged zebrafish Dram1 | This study |

## Notes

### Competing Interest Statement

The authors have declared no competing interest.

### Summary of Updates

- Author names updated - Location of figure labels changed to make them visible in the automatically generated pdf file

